# Seroprevalence and parasite rates of *Plasmodium malariae* in a high malaria transmission setting of southern Nigeria

**DOI:** 10.1101/725796

**Authors:** Eniyou C. Oriero, Adeola Y. Olukosi, Olabisi A. Oduwole, Abdoulaye Djimde, Umberto D’Alessandro, Martin M. Meremikwu, Alfred Amambua-Ngwa

**Author notes:** Corresponding author Medical Research Council Unit The Gambia at LSHTM, Fajara, P.O Box 273, Banjul, The Gambia, ext3016.

## Abstract

While *Plasmodium falciparum* continues to be the main target for malaria elimination, other *Plasmodium* species persist in Africa. Their clinical diagnosis is uncommon while rapid diagnostic tests (RDTs), the most widely used malaria diagnostic tool, are only able to distinguish between *P. falciparum* and non-falciparum species, the latter as ‘pan-species’. Blood samples, both from clinical cases and communites, were collected in southern Nigeria (Lagos and Calabar) in 2017 (October-December) and 2018 (October–November), and analysed by several methods, namely microscopy, quantitative real-time PCR (qPCR), and peptide serology targeting candidate antigens (*PmAMA1, PmLDH and PmCSP*). The sensitivity of the diagnostic approaches for the dominant *P. falciparum* were comparable, detecting approx. 80% infection, but not so for non-falciparum species – 3% and 10% infection for *P. malariae*; 0% and 3% infection for *P. ovale* by microscopy and qPCR respectively, across communities. *P. ovale* prevalence was less than 5%. Infection rates for *P. malariae* varied between age groups, with the highest rates in individuals > 5 years. *P. malariae* specific seroprevalence rates fluctuated in those < 10 years but generally reached the peak around 20 years of age for all peptides. The heterogeneity and rates of these non-falciparum species calls for increased specific diagnosis and targeting by elimination strategies.

## Introduction

Current global efforts towards malaria elimination target the dominant malaria parasites, *Plasmodium falciparum* and *Plasmodium vivax*.^1^ However, there are three other malaria species infecting humans, with *Plasmodium malariae* being the most common worldwide, including sub-Saharan Africa (sSA).^2^ *P. malariae* causes the so-called quartan malaria as the onset of fever occurs in an interval of three to four days, due to its 72-hour blood stage life cycle.^3,4^ Compared to falciparum malaria, it is a relatively benign infection, possibly due to low parasite densities, although it can cause severe anemia^5^ and nephrotic syndrome in both children and adults.^6,7^ *P. malariae* often is found in co-infections with other *Plasmodium* species,^3^ and because of this and the focus on *P. falciparum*, its biology, prevalence and specific public health impact have mostly been understudied.

Species-specific diagnosis of *P. malariae* is uncommon in routine malaria diagnosis; rapid diagnostic tests (RDTs), the most widely used diagnostic tools, predominantly target *P. falciparum* while other *Plasmodium* species are identified as ‘pan-species’.^8^ As a result, its burden and the population at risk in most endemic areas remain largely unknown. A recent study reported persistent transmission of *P. malariae* over a 22-year period in Tanzania, suggesting that the decline in *P. falciparum* prevalence could provide a favorable ecological niche for other malaria parasite species.^9^ This has also recently been reported for *P. knowlesi*, a zoonotic malaria species, in Malaysia.^10^

There were an estimated 228 million cases of malaria worldwide in 2018, 93% in Africa, with Nigeria contributing approximately 25% of these.^1^ Malaria transmission in Nigeria is perennial, mostly due to conducive geographic landscape, high temperatures and rainfall; about 85% of the population lives in areas of mesoendemic transmission.^11^ Although *P. falciparum* is the dominant species, responsible for over 95% of cases of clinical malaria, *P. malariae* is found in 9.8% of malaria cases, mostly as mixed infection.^11^ Nevertheless, its prevalence, mostly determined by molecular methods, can be as high as 26%.^12^ Here we report the prevalence of *P. malariae* infections as determined by molecular methods and the sero-prevalence of antibodies against this malaria species in individuals attending health facilities in two geopolitical and climatic zones of southern Nigeria.

## Methods

### Study site and Plasmodium spp survey

A preliminary survey was carried out between October and December 2017 in two study sites in southern Nigeria - Lagos in the south-west and Calabar in the south-south geopolitical zones, respectively (**Figure 1A**). Lagos is the most populous city in Africa with a population of approximately 15 million people; it has a tropical savanna climate, with a wet season between April and October, and a dry season in the following months. The wettest month is June, with mean precipitation of 315.5 millimetres (12.42 in). Located near the equator, Lagos has minimal seasonal temperature variation, with high temperatures ranging between 28.3–32.9 °C.^13^ Calabar has a tropical monsoon climate, with a long wet season (February to November) and a short dry season covering the remaining two months. Temperatures are relatively constant throughout the year, with average high temperatures ranging between 25 and 28°C; relative humidity is high with most months of the year recording a monthly mean value of 80%, and annual average precipitation just over 3,000 millimetres (120 in).^14,15^

**Figure 1:**
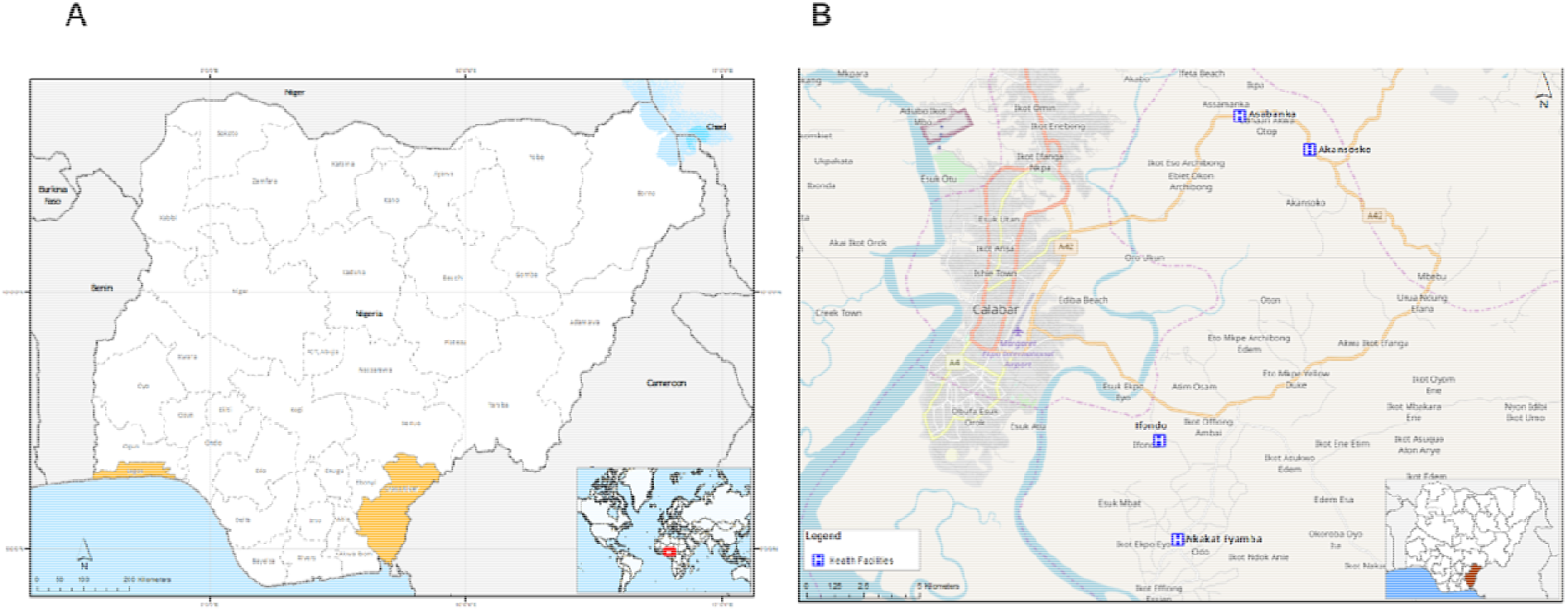
Map of Study sites in Nigeria showing A) Lagos (southwest) and Cross River (southeast) States; and B) Health facilities in Cross River State, where participants were recruited.

Suspected malaria cases at selected health facilities in both Lagos and Calabar were screened for malaria with a pan-species rapid diagnostic test (RDT) - SD Bioline, Cat No 05FK60 after providing a written informed consent to participate in the study. A blood sample was collected systematically by fingerprick for a thick blood film and dried blood spots (DBS) for molecular detection of *Plasmodium* species. At the peak of the following malaria transmission season (November/December 2018), a second survey was carried out in Akpabuyo local government area (LGA) of Cross River State, approximately 30 km from Calabar Municipality (**Figure 1B**). Participants who reported at health facilities were recruited and study sample was collected from them as well as RDT testing, following prior community sensitization.

### Ethical considerations

Ethical approval was obtained from the respective authorities in Lagos and Cross River State, References – IRB/17/038, CRSMOH/RP/REC/2017/545 and CRSMOH/RP/REC/2017/809. Written informed consent was obtained for adult participants while parental or guardian consent was obtained for minors. Participants did not receive any retribution for their participation in study. All individuals diagnosed with malaria by RDT were treated accordingly. Febrile individuals with a negative RDT were referred to the health facility staff for further investigation.

### Detection of Plasmodium spp parasites

Thick blood microscopy slides stained with 10% Giemsa solution were examined across 100 high power fields by two microscopists and parasite counts recorded per number of white blood cells observed. In case of≥ 20% variance in counts between primary and secondary microscopists, the counts closest to that of a third reader was retained. A slide was declared negative only after observing no parasite in 100 microscopic high-power fields while parasite density was calculated using WHO protocols; number of parasitized erythrocytes against 8000 white blood cells (WBCs),^16^ expressed as parasitaemia per microlitre of blood.

For molecular diagnosis of *Plasmodium spp*, DNA was extracted from DBS samples using the QIAamp DNA mini kit (QIAGEN, Germany) according to manufacturers’ protocols. For the preliminary survey, presence of *Plasmodium* species was detected using a commercial species-specific quantitative real-time PCR (qPCR) assay, Genesig® *Plasmodium* species kits (Primerdesign™ Ltd, UK) according to manufacturers’ protocol. Subsequently, presence of *P. malariae* and *P. ovale* were detected using a custom qPCR assay targeting the *Plasmepsin* gene,^17^ which showed comparable results with the Genesig® kit, while presence of *P. vivax* was not assessed further. A preamplification was done using 0.1 μM of the forward and reverse primers for *P. malariae* (CCWR2K1_Fwd TTCAGTCAGGATATGTAAAACAAAATTATTTAGGTA; CCWR2K1_Rev CCTACTTCCCCTTCACCATAAAACA) and *P. ovale* (CCRR9V5_Fwd

ACTCTTGGTTATTTGTCTGCACCTT; CCRR9V5_Rev CTATGTTACCATAAACAGGTTCTAAATCATCTGT), 1× Qiagen Multiplex Master Mix (QIAGEN, Germany) and 5 μL of template DNA in a 15 μL reaction. The main amplification reaction contained 3 μL of pre-amplified products in a 15 μL total reaction volume and was done using 1× qPCR Taqman Universal Master Mix (ThermoFischer Scientific, USA), 0.4 μM of the forward and reverse primers, and 0.2 μM of the labelled probe (*P. malariae* - CCWR2K1_Probe TCGTCTAGTTCTATTACGTCATTTTC and *P. ovale* - CCRR9V5_Probe TCAGTTGCTTCAACAAATTT). Preamplification was performed under the following conditions: initial denaturation for 5 min at 95°C, 12 cycles of denaturation for 30 s at 95°C, annealing at 60°C for 1 min, and extension for 90 s at 72°C; while the main amplification and detection were performed under the following conditions: initial denaturation for 10 min at 95°C, 40 cycles of denaturation for 10 s at 95°C, and 1 min at 60°C with data collection. *P. falciparum* was detected using a highly sensitive qPCR assay (varATS qPCR) targeting multi-copy subtelomeric sequences.^18^ Amplification reaction was done using 1× qPCR Taqman Universal Master Mix (ThermoFischer Scientific, USA).

### Peptide design and serology assays

*P. malariae*-specific peptides were designed from highly immunogenic epitopes of three *P. malariae* antigens – apical membrane antigen (AMA1), circumsporozoite surface protein (CSP), and lactose dehydrogenase (LDH) (**Table 1**). A *P. falciparum*-specific peptide was also designed from the same region as one of the PmAMA1 peptides to explore species specificity. A web tool, SVMTriP^19^ was used to predict linear antigenic epitopes from the whole *P. malariae* antigen sequences and the petides were queried for species specificity using NCBI BLASTP 2.9.0+. The regions of immunogenic epitopes containing the peptides were then compared with ortholgs of predicted *P. falciparum* epitopes available on the *Plasmodium* Database (PlasmoDB)^20^ and the Immune Epitope Database and Resource Analaysis (IEDB).^21^ The peptides indicating high species specificity and high immunogenicity were chosen for synthesis (thinkpeptides® ProImmune Ltd, Oxford, UK) at 95% purity and 1 - 4 mg synthesis scale.

**Table 1:**
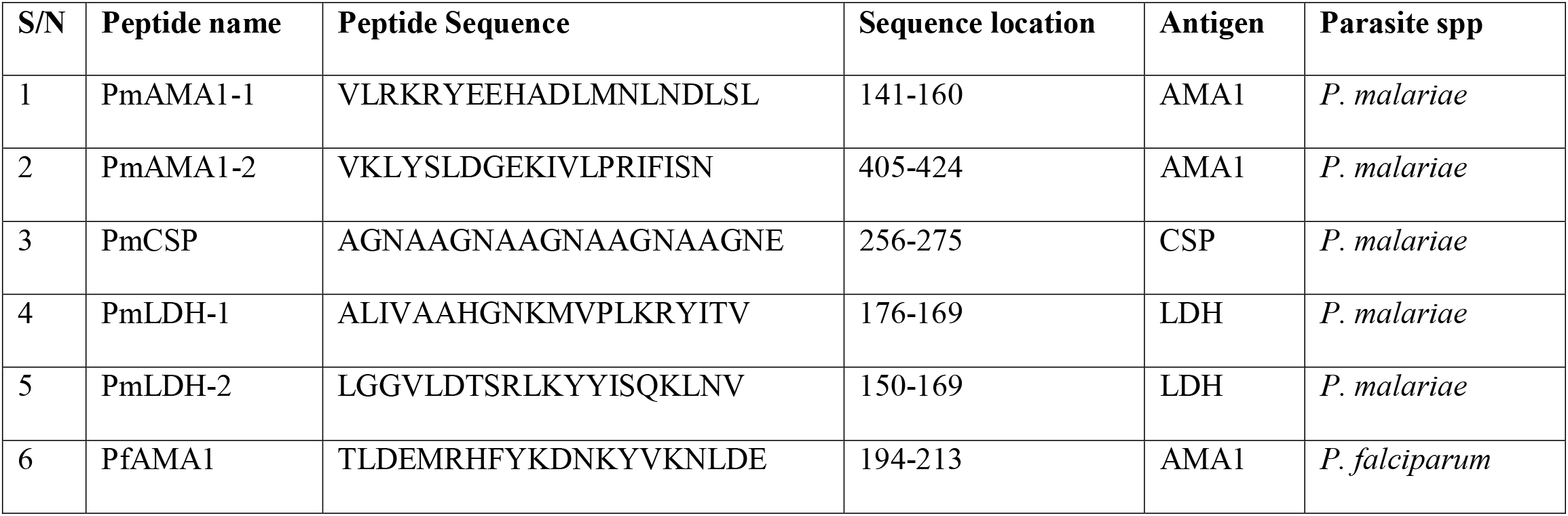
List of *Plasmodium spp* peptides for the serology assays

Antibodies against the *Plasmodium spp* peptides were detected by indirect ELISA on eluted filter paper samples. One disc (6 mm) per sample was punched and placed at the bottom of 96 well plates - flat bottom with low evaporation lids (Corning Inc. USA). Serum from dried blood spots (DBS) was eluted following overnight (18 h) incubation at 4°C in 150 μL of reconstitution buffer (1x Phosphate Buffer Saline; 0.2% Tween20). The reconstituted solution was equivalent to a 1:100 dilution of whole blood, corresponding to 1∶200 dilution of plasma or serum antibodies, if the blood were at 50% haematocrit.^22^ Peptides were coated on 96-well Nunc MaxiSorp™ Flat-Bottom Plates (Thermo Fischer, USA) at a concentration of 3 μg/mL. Following blocking of all remaing binding surfaces with 1% skimmed milk, DBS eluate was added at a final concentration of 1:2000 with respect to the corresponding plasma sample and analysed in duplicate wells per sample. Each ELISA plate included a positive control, which was a pool of DBS eluate from samples confirmed to be *P. malariae* positive by microscopy and qPCR. Also included in each plate were plate blanks (no serum) and negative control, which was un-exposed European pooled serum.^23^ Secondary detection was with Human IgG (H&L) Antibody Peroxidase Conjugated (Sigma – Aldrich, Germany), developed with Pierce™ TMB Substrate Kit (Thermo Fischer, USA). The reaction was stopped using 50 μl of Stop Solution for TMB Substrates (Thermo Fischer, USA). Optical densities (ODs) were read on the EMax® Plus microplate reader (Molecular Devices, USA) at 450 nanometer (nm).

### Data analysis

Data analysis was conducted using Microsoft Excel spreadsheet, GraphPad Prism Version 8.4.2 (La Jolla, California, USA), and Stata IC 16 (Stata Corp, College Station, TX, USA). Cross tabulations and t-test, Pearson Chi squared test (^2^) or Fisher’s exact test were used, where appropriate, to explore associations between *P. malariae* infection and demographic variables of interest. Analytical procedures for reducing between-plate variation included calculating the differences between the mean ODs averaged across the duplicates of the pooled positive control for each plate and the overall mean OD values across all plates; and obtaining the estimated plate effect for a specific plate by averaging the calculated differences across all pooled controls. Test samples on a given plate were then adjusted by subtracting the estimated plate effect from their OD values.^24^ Positivity for antimalarial peptide response was obtained by modelling their normalized FOC values (log2 transformed), using finite mixture models, Gaussian distribution with two components.^22,23^ The estimated mean of the narrower distribution was used as the mean for negative samples. The cut-off for seropositivity was obtained by taking the difference between the peptide FOC values and the mean of the narrower distribution plus 2 standard deviations.

Seroprevalence rates were analysed according the following age groups: 10 - 20 years, 20 – 30 years and > 30 years so there would be a similar number of individuals in each group. A three exponent non-linear regression model and line of best fit, using the quadratic estimation, was used to obtain the predicted probability of being seropositive for a particular peptide.

## Results

### Summary of participants recruited

In 2017, a total of 243 individuals with a positive malaria RDT were recruited, most of them in Calabar (71%, 172/243); children 6-10 years old were the largest age group (36%, 88/243) (**Table 2**). The following year, a total of 798 participants were recruited from four villages in Akpabuyo local government area, although the large majority (77%, 613/798) were from 2 villages (**Table 2**). The age distribution of the participants was different from that of the preliminary survey done in 2017; adolescents and adults represented about half of the participants (51%, 410/798), followed by children 11-15 years old (21%, 168/798) (**Table 2**).

**Table 2:**
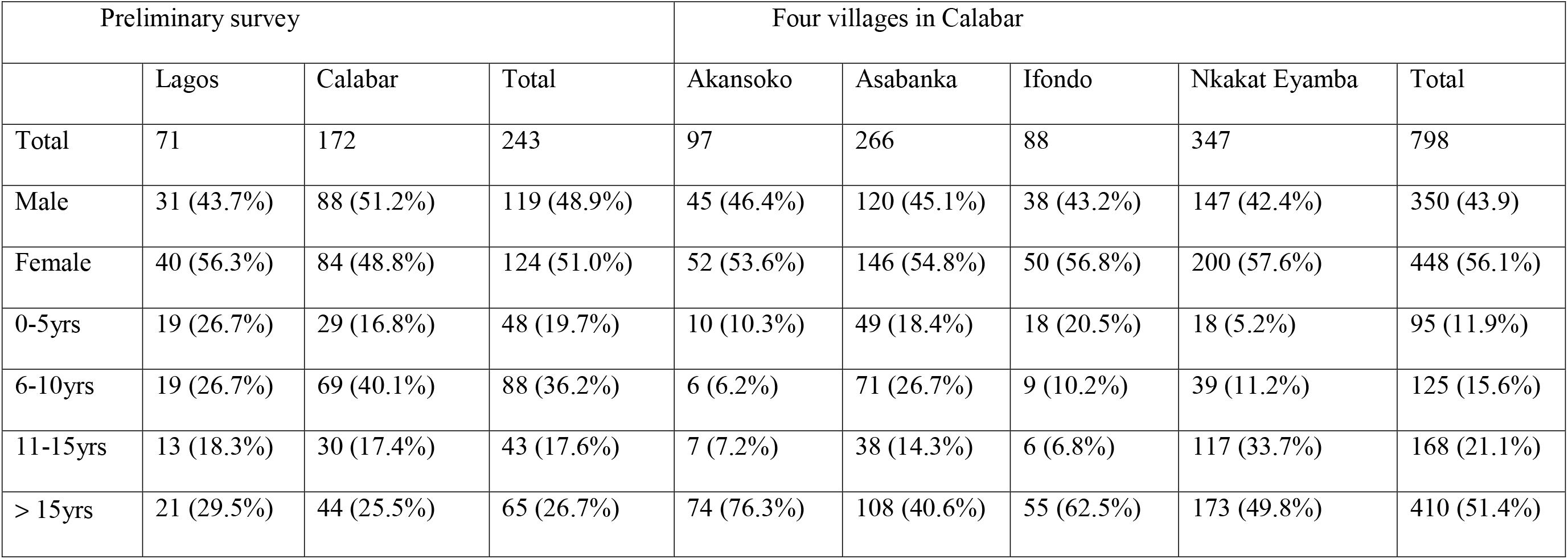
Summary of participants recruited in the two malaria transmission seasons

| Preliminary survey |  |  |  | Four villages in Calabar |  |  |  |  |
| --- | --- | --- | --- | --- | --- | --- | --- | --- |
|  | Lagos | Calabar | Total | Akansoko | Asabanka | Ifondo | Nkakat Eyamba | Total |
| Total | 71 | 172 | 243 | 97 | 266 | 88 | 347 | 798 |
| Male | 31 (43.7%) | 88 (51.2%) | 119 (48.9%) | 45 (46.4%) | 120 (45.1%) | 38 (43.2%) | 147 (42.4%) | 350 (43.9) |
| Female | 40 (56.3%) | 84 (48.8%) | 124 (51.0%) | 52 (53.6%) | 146 (54.8%) | 50 (56.8%) | 200 (57.6%) | 448 (56.1%) |
| 0-5yrs | 19 (26.7%) | 29 (16.8%) | 48 (19.7%) | 10 (10.3%) | 49 (18.4%) | 18 (20.5%) | 18 (5.2%) | 95 (11.9%) |
| 6-10yrs | 19 (26.7%) | 69 (40.1%) | 88 (36.2%) | 6 (6.2%) | 71 (26.7%) | 9 (10.2%) | 39 (11.2%) | 125 (15.6%) |
| 11-15yrs | 13 (18.3%) | 30 (17.4%) | 43 (17.6%) | 7 (7.2%) | 38 (14.3%) | 6 (6.8%) | 117 (33.7%) | 168 (21.1%) |
| > 15yrs | 21 (29.5%) | 44 (25.5%) | 65 (26.7%) | 74 (76.3%) | 108 (40.6%) | 55 (62.5%) | 173 (49.8%) | 410 (51.4%) |

### Molecular prevalence of Plasmodium species

In 2017, when only RDT positive individuals were recruited, the proportion of *Plasmodium* species detected in Lagos by qPCR were 81.7%, 1.4%, 1.4% and 0% for *P. falciparum, P. malariae, P. ovale*, respectively; whereas in Calabar, the proportions detected were 80.8%, 9.9%, 5.8%, respectively (**Figure 2**); no *P. vivax* infection was detected. There was no statistically significant difference between *P. falciparum* detected by qPCR and microscopy in the different study areas. However, significantly more *P. malariae* and *P. ovale* infections were detected by qPCR in Calabar compared to Lagos, emphasizing the low specificity of species specific characterisation of malaria parasites by microscopy. **Table 3** summarises the prevalence of *Plasmodium* spp detected as mixed or mono infections in the four different villages in Calabar. The two villages situated to the North of Akpabuyo LGA had significantly more non-falciparum species than the two villages to the South (**Figure 3**). Despite observing a higher proportion of *P. malariae* infections in older individuals (> 15 years), the odds of having *P. malariae* as mono infection was the highest in the < 5 years age group. The least odds for monoinfections were in those of ages 11 - 15 years (**Table 4**); the odds of *P. malariae* mono infections were the highest in Asabanka, with 4.5% mono infection, and the least in Ifondo community, with 1.1% mono infection (**Tables 3 and 4**).

**Table 3:**
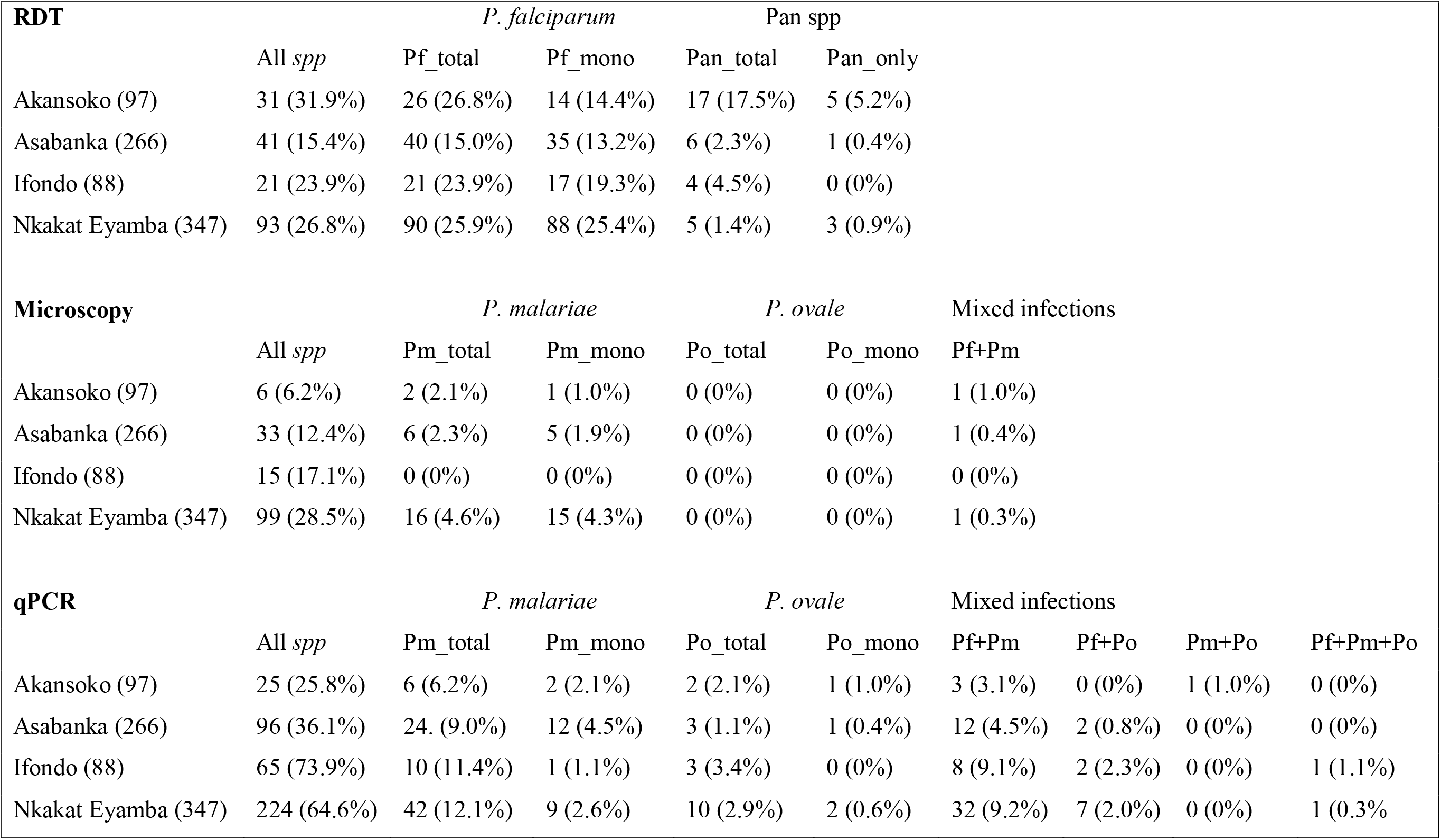
Summary of non-falciparum species detected in south-south Nigeria as mixed and mono infections by the different diagnostic methods – RDT, Microscopy and qPCR.

**Table 4:** Odds of *P. malariae* occurring as mono infections or mixed infections with other *Plasmodium spp*. in the four different villages and across age groups.

|  | Mono<br>infections | Mixed<br>infections | Total | Odds# | 95% CI | Odds<br>ratio* | 95% CI |
| --- | --- | --- | --- | --- | --- | --- | --- |
| Village |  |  |  |  |  |  |  |
| Akansoko | 2 | 4 | 6 | 0.5 | 0.09 - 2.73 | 4.5 | 0.25 - 80.57 |
| Asabanka | 12 | 12 | 24 | 1.0 | 0.45 - 2.23 | 9 | 0.79 - 101.81 |
| Ifondo | 1 | 9 | 10 | 0.11 | 0.01 - 0.88 | 1 | . . |
| Nkakat Eyamba | 9 | 33 | 42 | 0.27 | 0.13 - 0.57 | 2.46 | 0.26 - 22.82 |
| Age group |  |  |  |  |  |  |  |
| 0-5yrs | 4 | 3 | 7 | 1.33 | 0.29 - 5.96 | 16.67 | 1.39 - 199.42 |
| 6-10yrs | 3 | 9 | 12 | 0.33 | 0.09 - 1.23 | 4.17 | 0.55 - 31.75 |
| 11-15yrs | 2 | 25 | 27 | 0.08 | 0.02 - 0.34 | 1 | . . |
| >15yrs | 15 | 21 | 36 | 0.71 | 0.37 - 1.39 | 8.93 | 1.58 - 50.32 |

#Odds of *P. malariae* observed as a mono infection compared to mixed infections
\*Odds ratio calculated relative to the least odds

**Figure 2:**
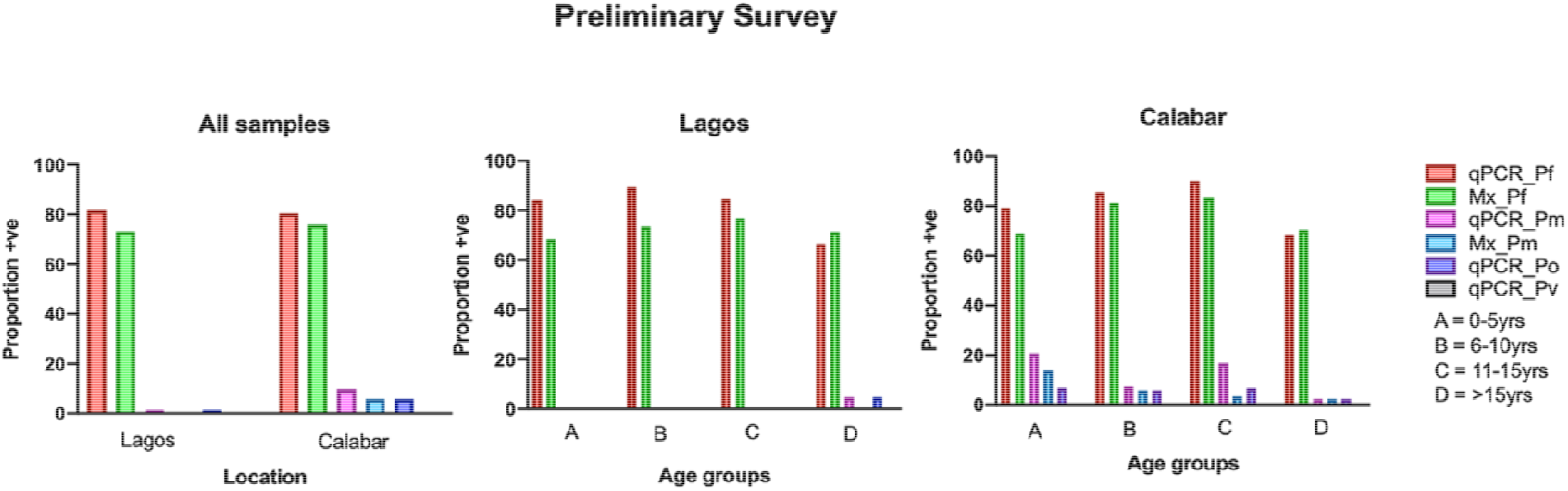
Prevalence of *Plasmodium* species in RDT positive individuals from the preliminary survey. qPCR_Pf = *P. falciparum* detected by qPCR; Mx_Pf = *P. falciparum* detected by Microscopy; qPCR_Pm = *P. malariae* detected by qPCR; Mx_Pm = *P. malariae* detected by Microscopy; qPCR_Po = *P. ovale* detected by qPCR and qPCR_Pv = *P. vivax* detected by qPCR.

**Figure 3:**
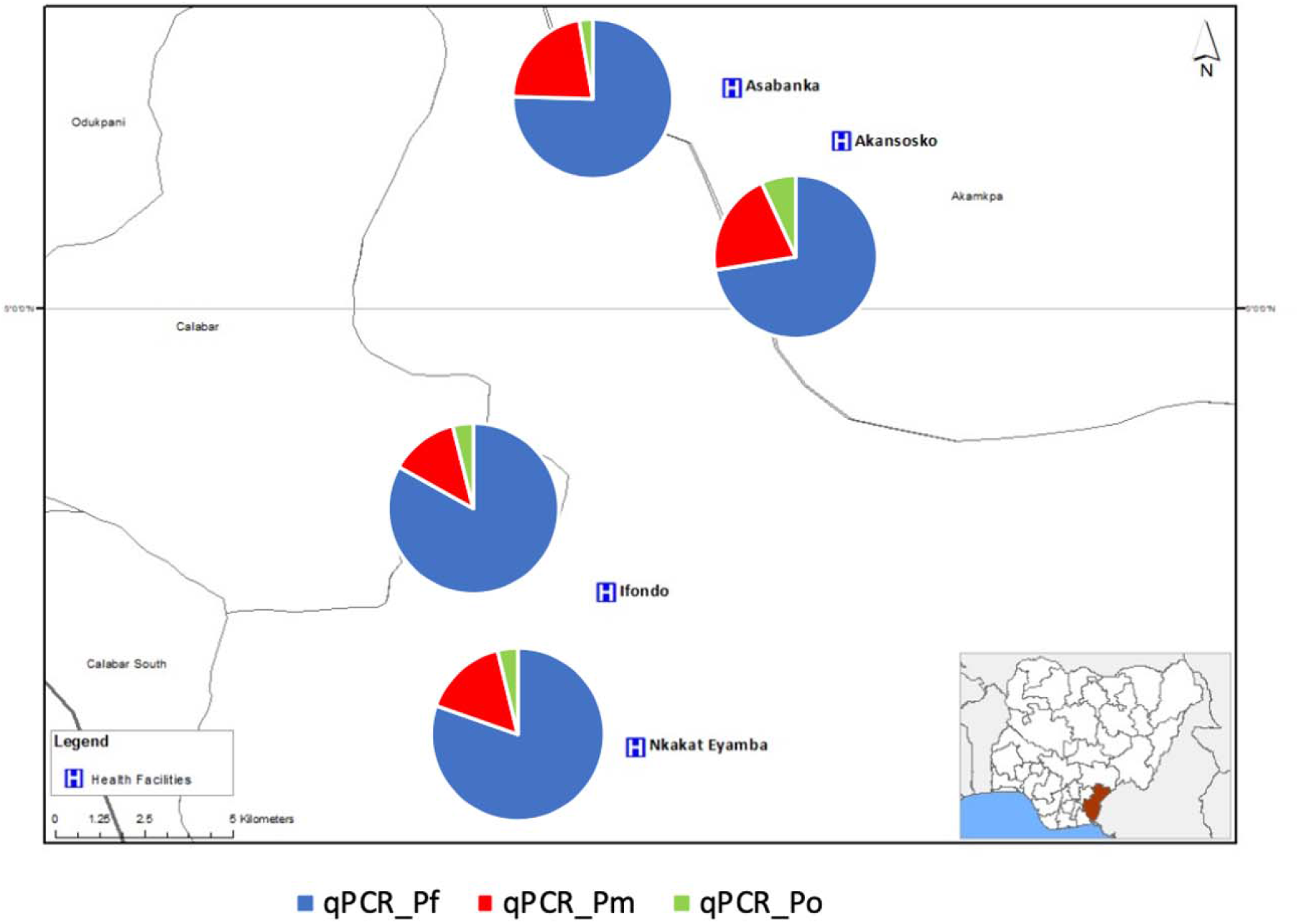
Proportion of *Plasmodium* species detected in the four villages in southeastern Nigeria, detected by qPCR (qPCR_Pf = *P. falciparum*, qPCR_Pm = *P. malariae*, qPCR_Po = *P. ovale*)

### Serological responses to the different P. malariae peptides

Anti-peptide IgG antibody levels were detected against each of the *Plasmodium spp* peptides designed and these were relatively higher than the median values of a pool of non-immune sera, reported as fold over the negative control (FOC). Over 70% of the samples showed antibody reactivity levels above the negative control for each of the peptides (PmAMA1-1 = 73.1%, PmAMA1-2 = 86.7%, PmCSP = 78.2%, PmLDH-1 = 80.9%, PmLDH-2 = 87.2% and PfAMA1 = 80.3%). From the area under the Receiver Operator Curve (ROC) for the different peptides, the FOC responses of all the peptides could predict up to 70% of the results obtained by microscopy (range = 62.6% - 70.5%) and RDT (range = 66.4% - 71.9%) (**Figure 4**). The *P. malariae*-specific anti-peptide antibody reactivities relative to the *P. falciparum* peptide for samples from the preliminary survey expressed significantly higher *P. malariae* responses in Calabar compared to Lagos with the exception of PmCSP peptide to which Lagos samples were more reactive (**Supplementary figure 1**). Due to the small number of samples from Lagos and some of the villages in Calabar, geographical location was not considered in all subsequent analysis. As observed with *P. falciparum*, seroprevalence increased with age for all the *P. malariae* peptides from the age of 10 years (**Figure 5**). Proportions of seropositivity in age groups below 10 years were variable between antigens, with no consistent pattern between peptides (**Supplementary figure 2**). The proportion of seropositivity decreased for PmLDH-1, while it increased for PfAMA1 for this younger age group.

**Figure 4:**
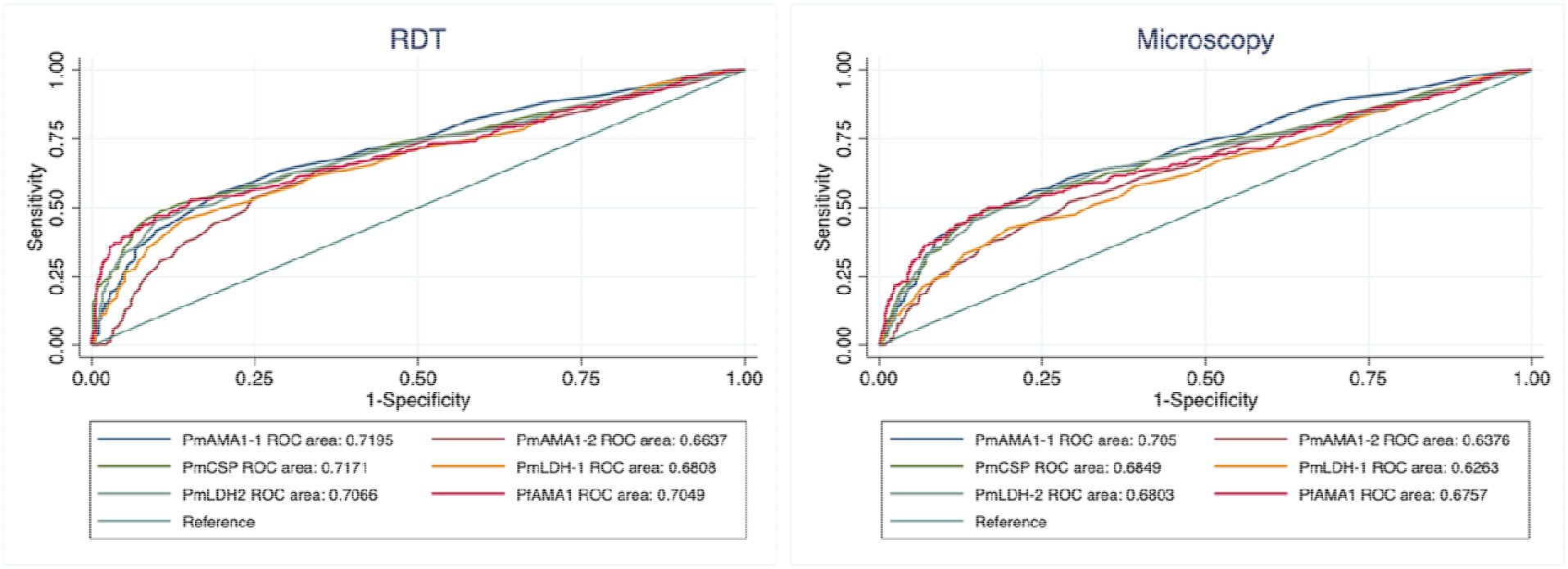
Area under the receiver operator curve (ROC) showing comparison of fold over the negative control (FOC) for each of the peptides with RDT and microscopy results.

**Figure 5:**
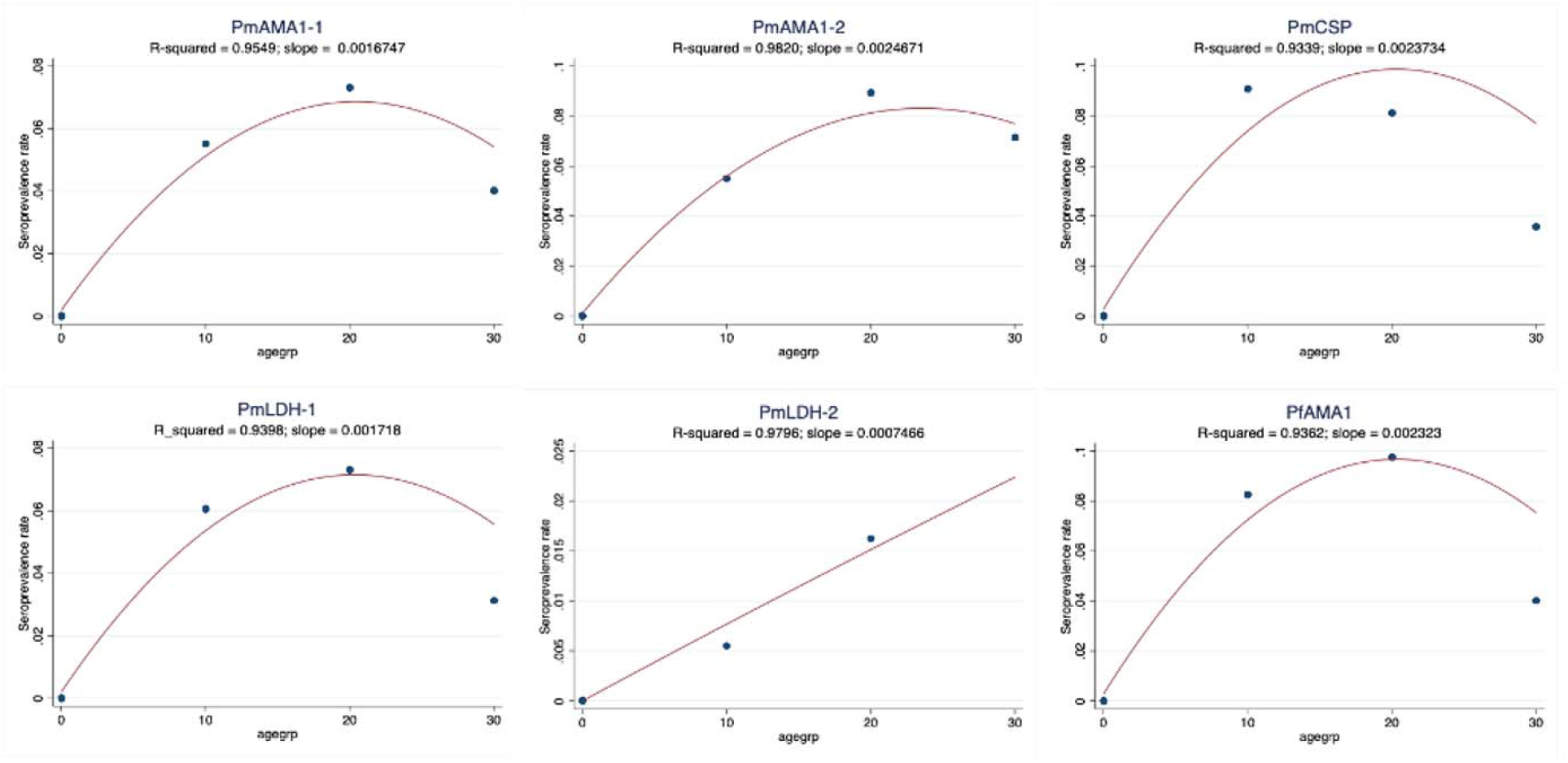
Seroprevalence rates of the different species-specific peptides in different age groups, the slope is the estimated change of rate over age group.

## Discussion

Prevalence of non-falciparum species was extremely heterogeneous in communities in the western and eastern regions of southern Nigeria supporting the need for broadening the focus of the National Malaria Control Program beyond *P. falciparum*.^25–28^ Nigeria accounts for an estimated 25% of global malaria cases, mostly considered to be due to *P. falciparum* infection.^29^ With the push for malaria eradication, this indicates that other human *Plasmodium* species should also be considered in current elimination strategies and evaluation of intervention outcomes. Unlike *P. falciparum*, *P. malariae* infections were more prevalent in individuals older than 5 years. Current malaria intervention strategies such as seasonal malaria chemoprevention (SMC) usually target children 3 - 59 months old. Hence older age groups in *P. malariae* endemic areas could serve as a reservoir for future expansion of this parasite if *P. falciparum* is selectively eliminated. Higher prevalence of *P. malariae* infections were also observed in older age groups in Papua New Guinea based on a sensitive molecular diagnostic method, post-PCR ligase detection reaction-fluorescent microsphere assay (LDR-FMA), compared to light microscopy.^30^ These comparable results despite the wide geographic distance between Nigeria and Papua calls for wider surveillance and reporting of these “minor” malaria parasite species and broadening of malaria elimination strategies to include all population age groups at risk of infection with *P. fal*ciparum and non-falciparum species.

Despite the lower prevalence of infection in children < 5 years of age, the odds of *P. malariae* mono infections were higher in this age group than in individuals aged > 15 years. This could reflect the level of exposure and immunity, as younger children may be protected by maternal antibodies as well as interventions such as insecticide treated bed nets and SMC.^31^ The age distribution also mirrors the pattern seen for *P. falciparum*, where an age shift in high infection rates in older children has been reported across Africa following large scale up of malaria interventions.^32^ The odds of *P. malariae* mono infection also varied between communities, with lowest infection odds in Ifondo community, which had, by molecular methods, the highest prevalence of *P. falciparum* infections. This needs further investigation due to the small numbers reported here, reflected in the wide confidence intervals. Nonetheless, some studies have reported higher *P. malariae* prevalence rates when outside the *P. falciparum* peak transmission, suggesting an interaction between the occurrence of these species.^33–35^ Such dynamics could be driven by co-immune mechanisms as the genomes and proteins of these species are highly similar,^36^ hypothetically allowing for high immune responses against *P. falciparum* to eliminate *P. malariae* from co-infections.

There could also be an ecological dynamics in which higher multiplication rates and shorter cycle time of *P. falciparum* would displace *P. malariae*, which often occurs at low parasite densities and prefer to invade older red blood cells.^4^ Infected RBCs are cleared by the spleen during malaria infections,^37,38^ which may in turn remove *P. malariae* from peripheral circulation. However, *P. malariae* has been reported persist following treatment of mixed species malaria with ACTs,^39–41^ an indication that further studies are needed to elucidate the dynamics and consequences of malaria parasite co-infections. This will require specific surveillance tools, including sensitive and specific diagnostic and serological assays.

In the absence of whole antigen ELISA assays for *P. malariae* and other non-falciparum infections, species-specific peptide ELISA assays provided an alternative tool for determining seroprevalence of antibodies against *P. malariae*. Predicted peptide epitopes from *P. malariae* AMA1, CSP, and LDH proteins were all recognised by human IgG antibodies, evidence of specific immune responses in exposed communities. Higher relative antibodies were seen in the South-Eastern region with higher malaria transmission. Thus *P. malariae* elicits specific antibody responses, which could be targeted for the development of sensitive surveillance tools. Seroprevalence has been used as a surrogate for exposure, enabling the determination of force of infection (FOI) and age specific immunity for *P. falciparum* and *P.vivax*, in endemic regions.^23,42–44^ ELISA techniques for detection of antibodies have also been widely used for the diagnosis of infectious diseases and have been used in malaria since the 1970s.^45^ Adequately powered studies to compare and predict transmission intensity of *P. malariae* and other non-falciparum species across sub-Saharan Africa would be useful for strategizing towards control and elimination of all malaria parasite species.

The preliminary survey carried out in 2017 showed a lower prevalence of non-falciparum species in Lagos compared to Calabar. This was not surprising as urbanization in Lagos metropolitan city, with better housing and access to health services, will limit malaria transmission despite similar vector populations.^46^ A consistent pattern of increased urbanization coincident with decreasing malaria transmission and elimination over the past century has been reported.^47^ The non-falciparum infections in the urban area were submicroscopic and could only be detected by sensitive molecular methods. This emphasizes the challenges to malaria diagnosis with decreased prevalence and more sub-microscopic infections. Sensitive and accessible diagnostic tools for routine detection of all human *Plasmodium* species therefore remain important. Microscopy, which is still considered the gold standard for malaria diagnosis, is laborious and time consuming for *Plasmodium* speciation, requiring expert microscopists.

Current antigen-based pan-species RDT also present peculiar challenges, including low sensitivity and non-species specificity.^48^ The poor sensitivity for pan-species *Plasmodium* detection was recently demonstrated in an experimental human blood-stage model for *P. malariae* infection, where RDT targeting pan-genus lactate dehydrogenase enzyme – pLDH, remained negative despite the presence of symptoms consistent with malaria 72-hour prior to testing.^49^

Most of the malaria infections detected by PCR were low-grade infections or sub-microscopic, which is a well documented limitation of microscopy and this may explain the high discrepancy between *Plasmodium* species detected by microscopy compared with PCR. As for *P. falciparum*, ultra-sensitive molecular diagnostics that target multicopy genes will be relevant to assess the true burden of non-falciparum malaria infections.

Despite recent reports of *P. vivax* detection in Duffy-negative individuals in sub-Saharan Africa, including South Western Nigeria,^50^ no *P. vivax* infection was detected. The prevalence of *P. vivax* in Duffy negative individuals could therefore be specific to some susceptible populations driven by yet undetermined biological and environmental factors.

In conclusion, the prevalence rates of non-falciparum species reported here indicates that approaches for routine diagnosis in endemic settings should be encouraged to ascertain the true burden of these species as well as for appropriate interventions towards pan-species malaria elimination. Studies to determine the prevalence and dynamics of non-falciparum species as well as associated risk factors in endemic populations need to be expanded across sub-Saharan Africa.

## Supporting information

Supplementary docs

## Acknowledgment

We would like to acknowledge Ernest Oriero for designing and adapting all maps used, Bekai Njie for the primary microscopy reads, Dr Sola Ajibaye for helping out with logistics in the Lagos site and the Statistics Department of the Medical Research Council Unit The Gambia at LSHTM for support with the data analysis.

## Financial Support

This work was supported through the DELTAS Africa Initiative [DELGEME grant 107740/Z/15/Z]. The DELTAS Africa Initiative is an independent funding scheme of the African Academy of Sciences (AAS)’s Alliance for Accelerating Excellence in Science in Africa (AESA) and supported by the New Partnership for Africa’s Development Planning and Coordinating Agency (NEPAD Agency) with funding from the Wellcome Trust [DELGEME grant 107740/Z/15/Z] and the UK government.

## Disclosures

The authors have no competing interests. The views expressed in this publication are those of the author(s) and not necessarily those of AAS, NEPAD Agency, Wellcome Trust or the UK government.

## Author Contributions

ECO conceptualized the study and is the principal investigator overseeing all aspects of the project, contributing to data curation, formal analysis and manuscript writing; AYO coordinated aspects of the investigation in the sites in Lagos; OO coordinated aspects of the investigation in the sites in Calabar; UDA and MM contributed to review and editing of the manuscript; AD contributed to funding of the research project and reviewing the manuscript; AAN contributed to conceptualization and supervision of the study, review and editing of the manuscript.

## Notes

### Competing Interest Statement

The authors have declared no competing interest.

### Summary of Updates

Seroprevalence data for P. malariae has been added to complement the molecular prevalence data.

