## Supplementary docs for "Seroprevalence and parasite rates of *Plasmodium malariae* in a high malaria transmission setting of southern Nigeria": Suppl.docx

**Supplementary document**

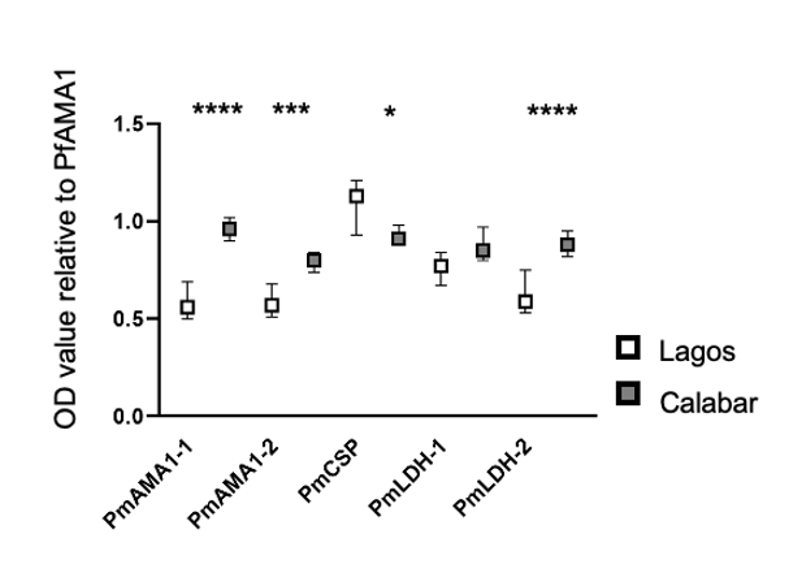

**Supplementary figure 1**: Comparison of malaria responses from the two study sites (preliminary survey) to the different *P. malariae* peptides, with respect to the response of the *P. falciparum* peptide.

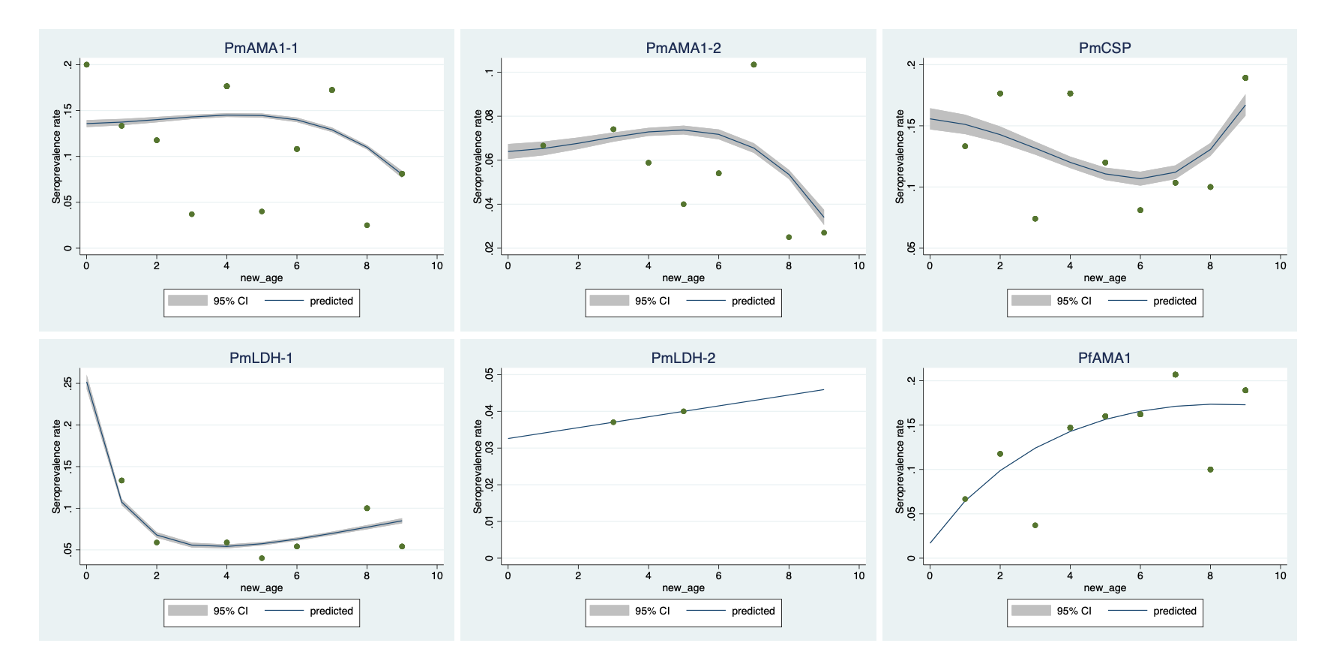

**Supplementary figure 2**: Seroprevalence rates of the different species-specific peptides in ages < 1 years to 9 years old.
